# The Cyclophilin Inhibitor Rencofilstat Decreases HCV-induced Hepatocellular Carcinoma Independently of Its Antiviral Activity

**DOI:** 10.1101/2023.08.19.553982

**Authors:** Winston Stauffer, Michael Bobardt, Daren Ure, Robert Foster, Philippe Gallay

**Author notes:** Corresponding authors. Philippe A. Gallay: Department of Immunology & Microbial Science, IMM-9, The Scripps Research Institute, 10550 N. Torrey Pines Rd., La Jolla, CA 92037.

## Abstract

There is an urgent need for the identification of new drugs that inhibit HCV-induced hepatocellular carcinoma (HCC). Our work demonstrates that cyclophilin inhibitors (CypI) represent such new drugs. We demonstrated that the non-immunosuppressive cyclosporine A (CsA) analog (CsAa) rencofilstat possesses dual therapeutic activities for the treatment of HCV infection and HCV-induced HCC. Specifically, we showed that HCV infection of humanized mice results in the progressive development of HCC. This was true for four genotypes tested (1 to 4). Remarkably, we demonstrated that rencofilstat inhibits the development of HCV-induced HCC in mice even when added 16 weeks post-infection when HCC is well established. Importantly, we showed that rencofilstat drastically reduces HCC progression independently of its anti-HCV activity. Indeed, the CypI rencofilstat inhibits HCC while other anti-HCV agents such as NS5A (NS5Ai) and NS5B (NS5Bi) fail to reduce HCC. In conclusion, this study shows for the first time that the CypI rencofilstat represents a potent therapeutic agent for the treatment of HCV-induced HCC.

## Introduction

The name, “cyclophilin” (Cyp), comes from the discovery of cyclophilin A (CypA) as a ligand of cyclosporine A (CsA) (1) and its peptidyl-prolyl isomerase (PPIase) activity (2). Cyps belong to a family of enzymes, which catalyzes the *cis-trans* isomerization of proline peptide bonds, which are unique from other bonds due to their ability to switch between *cis* and *trans* conformations. The Cyps-mediated catalyzation of *cis-trans* proline peptide bonds occurs, at many orders, faster than uncatalyzed *cis-trans* proline peptide bonds (3–4). Prolines shape the configuration of proteins due to the rigidity of the pyrrolidine ring of prolines. Cyps, by catalyzing the *cis-trans* isomerization of proline peptide bonds, regulate the structural conformation of proteins, their ligand-binding properties and biological functions. Thus, Cyps participate in a broad range of activities including i) initial folding of nascent peptides into proteins; ii) restraint of protein aggregation; iii) intracellular protein trafficking and secretion; iv) amplification of the second messenger signaling; and v) regulation of protein-protein interactions (5–8). The immunosuppressant CsA came into medical use in 1983 to prevent graft-versus-host disease (9). The immunosuppressive activity of CsA originates from CsA binding to CypA and formation of a ternary complex with calcineurin, which blocks activation of nuclear factor of activated T cells and its downstream signaling. CsA neutralizes the isomerase activity of CypA and other Cyp members such as CypB, CypC and CypD with high affinities (Ki of ∼15 nM) due to their highly conserved enzymatic pockets (10). Chemical alteration of CsA resulted in the identification of CsA analogs (CsAa) devoid of immunosuppressive activity, but with preserved CypA-binding activities (11–13). Four Cyp inhibitors (CypI) have been studied in humans – the CsA analogs (CsAa) NIM811, alisporivir (ALV)/Debio-025, SCY-635 and rencofilstat (CRV431).. Rencofilstat potently inhibits all cyclophilin isoforms tested-A, B, D, and G (14). Inhibitory constant or IC_50_ values ranged from 1 to 7 nM, which was up to 13 times more potent than the parent compound, CsA from which rencofilstat was derived. Other rencofilstat advantages over CsA as a nontransplant drug candidate include significantly diminished immunosuppressive activity, less drug transporter inhibition, and reduced cytotoxicity potential (14). Oral dosing to mice and rats led to exposures expected to be in line with therapeutic blood concentrations, and a 5- to 15-fold accumulation of rencofilstat in liver compared with whole blood concentrations across a wide range of rencofilstat dosing levels (14). More recently, rencofilstat was safe and well tolerated after 28 days in subjects with presumed F2/F3 NASH (15). The presence of NASH did not alter its pharmacokinetics.

We and others showed that CsAa inhibit HCV *in vitro* (16–33). ALV showed high anti-HCV efficacy in humans in phase I-III studies (34–37). We reported that HCV fails to infect CypA-knockdown (KD) cells while it infects CypB-KD, CypD-KD and parental cells (26), suggesting that HCV requires CypA to optimally replicate in human hepatocytes. Supporting this notion, the Ploss lab demonstrated that HCV fails to infect CypA-knockout (KO) humanized mice (38). We showed that HCV infection of CypA-KD cells is restored after reintroduction of wild-type CypA, but not isomerase-deficient CypA (H126Q mutation in the enzymatic pocket of CypA) (26), further suggesting that HCV relies on the foldase activity of CypA to replicate in cells. We demonstrated that CsAa inhibit HCV infection by preventing NS5A-CypA interactions and preventing the formation of ER double membrane vesicles (DMVs) required to shield the amplification of the viral RNA genome (28). Thus, the requirement for CypA in HCV infection is well understood at a cellular level. Moreover, we demonstrated that rencofilstat inhibits HCV-induced HCC in humanized mice and that viral replication was totally suppressed after 4 days of rencofilstat treatment (39). Altogether these data strongly suggest that rencofilstat inhibits HCV infection by sequentially neutralizing the PPIase activity of CypA, preventing CypA-NS5A interactions, leading to the suppression of CypA-NS5A-mediated DMV formation that normally protects the viral genome replication and viral particle formation.

Anti-HCV treatments have progressed significantly during the last 20 years. Current HCV treatments include highly successful combinations of pangenotypic direct-acting antivirals (DAA) with short period treatments (8–12 weeks), high sustained virological response (SVR) (>95%), and minimal side effects. However, the combination of specific viral genotypes (GT) and disorder conditions diminishes SVR levels to DAA such as HCV genotype 3 (GT3) infection, cirrhosis, and DAA resistance associated with the selection of resistance-associated substitutions (RASs) present at baseline or are acquired during treatment (40). An option to avoid the retreatment of HCV patients for viral resistance and GT3 infection would be to incorporate into current FDA-approved DAA regimens, antivirals with high barrier to resistance and with different antiviral mechanisms of action (MoA). The CypI rencofilstat possesses such unique anti-viral activities.

HCC is the third major cause of cancer mortality, globally accounting for 800,000 deaths per year (41). Moreover, HCC is one of the fastest-growing causes of cancer mortality in the United States (41). Current treatment for HCC includes liver transplantation, segmentectomy, chemotherapy, and systemic drug therapy. However, tumor recurrence may occur after transplantation and segmentectomy, and HCC is minimally responsive to chemotherapy. Moreover, chemotherapy presents numerous toxic side effects (42–43). Importantly, unexpectedly high rates of HCC recurrence occur after hepatic resection (44) and chemotherapy (45–46). Therefore, the identification of new effective treatments for HCC is a crucial research interest. We recently demonstrated and reported that rencofilstat decreases liver fibrosis in a non-viral 6-week carbon tetrachloride model as well as in a mouse model of nonalcoholic steatohepatitis (NASH) (47). Rencofilstat administration during a late, oncogenic stage of the NASH disease model results in a 50% reduction in the number and size of liver tumors (47). These findings are consistent with rencofilstat targeting fibrosis and cancer via multiple Cyp-mediated mechanisms and re-emphasize the utmost importance of exploiting rencofilstat as a safe and effective drug candidate for the treatment of liver diseases.

Approximately 71 million people worldwide are infected with HCV (48–50). In HCV patients, the liver damage varies from negligible injuries to pronounced fibrosis, cirrhosis, or HCC. Worldwide deaths associated with HCV-induced complications are estimated to be ∼600,000 in 2017 (20). In this study, we investigated for the first time whether rencofilstat could inhibit HCC development induced by HCV infection rather than induced by chemicals or high fat diet. We found that a daily rencofilstat treatment of mice dramatically diminished HCV-induced HCC in humanized mice even when added months post-infection when viral replication is robust and when HCC is established, suggesting that rencofilstat represents a novel therapeutic drug for the treatment of HCV-induced HCC in HCV patients infected even for several months.

## Materials and methods

### Drugs

Rencofilstat (CRV431) was synthesized in-house by chemical modification of cyclosporin A and its purity >95% was determined by HPLC. The NS5B polymerase inhibitor (NS5Bi) sofosbuvir and the NS5A inhibitor velpatasvir (NS5Ai) were purchased from MedChemExpress.

### Animal care

#### Animal housing

individually ventilated cage (IVC) racks are used to house the majority of mice. HEPA-filtered air is supplied into each cage at a rate of 60 air changes per hour. Mice are housed in solid bottom cages. Static mouse cages are changed at least once a week. IVCs are changed at least once every 14 days. Certain strains of rodents (e.g., diabetic) are changed into clean cages more frequently as needed. Room environment: heating, ventilation and air conditioning performance is routinely assessed as part of facility renovations, system repairs, and at least once every 3 years. Each animal room is equipped with a high/low thermo-hygrometer and its own computerized controlled thermostat. Animal care staff monitor and record animal room high/low temperatures and humidity daily on the room activity log. Temperature settings are consistent with Guide recommendations and are calibrated by the Engineering Department. Alarm points are set at ± 4°F. High or low temperature alarms are annunciated to the engineer on duty 24 hours a day. The Department of Animal Resources (DAR) management is notified of excursions. Most of the animal facilities are also equipped with an Edstrom Industries Watchdog environmental monitoring system in addition to the automated building management system (BMS). The Watchdog system registers temperature and humidity and sends alarms to Animal Resources management personnel. Humidity levels are not controlled in any of the facilities but are reliably maintained between 30–70% most of the year.

#### Diet

Food (Teklad LM-485 autoclavable diet) is provided ad libitum to mice in wirebar lids. Water: our vivarium is equipped with a reverse osmosis (R/O) water purification system and automatic watering distribution system from Edstrom Industries. DAR receives monthly water quality reports from the City of San Diego. R/O purified water is monitored daily during the workweek. Parameters are monitored including conductivity, temperature, pH level and chlorine concentration. Automatic water delivery systems (room and rack distribution lines) are timed for daily in-line flushing. Quick disconnect drinking valves are sanitized with each cage change or more often if needed. System sanitation and preventive maintenance is performed by the DAR equipment technicians.

#### Acclimation period

mice are allowed up to 72 hours to stabilize into their new housing environment. Some experimental paradigms involve examining the behavioral response to novelty and therefore the animal cannot be habituated to the procedure.

#### Animal suffering

To minimize suffering, all surgical procedures are carried out under anesthesia using isoflurane (1-4%) in conjunction with ketamine/xylazine ip (90 mg/kg and 10 mg/kg). Mice are monitored every 15 minutes after induction for respiratory and heart rates if the surgical procedure requires more time. Animals are provided buprenorphine (0.05–2.5 mg/kg s.c.) for 6– 12 h followed by flunixine meglumine (2.5 mg/kg s.c.) as a postoperative analgesic for 2 days post-implantation. Mice are observed 2 h, 6 h and 24 h post-surgery with daily monitoring during the study. Mice are supplied with acidified water supplemented with sulfamethoxazole (or sulfadiazine) with trimethoprim at a final concentration of 0.65–1.6 mg/mL to reduce chances of opportunistic bacterial colonization. MUP-uPA-SCID/Beige mice were maintained at DAR at TSRI in accordance with protocols approved by the TSRI Ethics Committee, the Institutional Animal Care and Use Committee (Protocol Number: 11–0015). This study was carried out in strict accordance with the recommendations in the Guide for the Care and Use of Laboratory Animals of the National Institutes of Health. All efforts were made to minimize suffering. The method of sacrifice used for the experimental mice was cervical dislocation. A power calculation was used to determine the sample size (number of mice/group). Ten mice per group were used for each treatment in all experiments.

### HCV chimeric mouse study

Transgenic mice carrying the uPA gene driven by the major urinary protein promoter were crossed onto a SCID/Beige background (MUP-uPA-SCID/Beige) (51). These transgenic mice are healthier than former Alb-uPA mice and provide an extended window from age 4 to 12 months for engraftment with human hepatocytes and infection with serum from infected chimpanzees (100 infectious doses): HC-TN GT1a, HC-J6 GT2a, S52 GT3a and ED43 GT4a (gift from Dr. Lanford). MUP-uPA-SCID/Beige mice (gift from A. Kumar) (4 months old) were transplanted with human hepatocytes (10^7^ cells/mouse). We usually obtain ∼300-500×10^6^ hepatocytes per donor (gift from D. Geller) as described previously (39). In brief, fresh hepatocytes were transplanted immediately upon arrival within 12–16 hour after isolation. Viable cell counts were determined. One cm skin incision was made in the upper left quadrant of the mouse abdomen to visualize the spleen. Human hepatocytes were injected intra-splenically. Incision was then closed with Vetbond tissue adhesive (3M Animal Care Products, St. Paul, MN). To verify the degree of “humanization”, blood was weekly collected for human albumin quantification by ELISA (Bethyl Laboratories) according to the manufacturer’s protocol. Mice expressing >300 μg/mL of human albumin (hAlb) were randomized into groups of 10 mice. It is crucial to note that the expression of the uPA transgene causes damage to the murine liver that is rapidly replaced by clusters of implanted human hepatocytes that can be visualized by hAlb immunostaining (51). We determined the optimal number of human hepatocytes needed to achieve an adequate level of engraftment by increasing the number of cells used in the engraftment and assessed for the functionality of repopulated hepatocytes by measuring hAlb in the serum of the mice by ELISA 30 days after engraftment. While mice that were engrafted with a range of 0.5 x 10^6^ to 2.0 x 10^6^ hepatocytes expressed a mean of 320 ug/mL of hAlb, increasing the number of hepatocytes to 4-6 x 10^6^ cells per mouse significantly improved reconstitution and resulted in a mean of 1.9 mg/mL (***P= 0.0006) (51). MUP-uPA-SCID/Beige mice that had been engrafted with human hepatocytes were then infected intravenously (i.v.) with concentrated viruses (genotype 1a, 2a, 3a and 4a) derived from cell culture (39). CRV431 was dissolved in polyethylene glycol 300 molecular weight (PEG-300), Velpatasvir and sofosbuvir were dissolved in DMSO and subsequently in 95% sterile saline solution. Drugs were administered once by oral gavage at 50 mg/kg at the indicated time points. Blood was collected retro-orbitally at the indicated time points.

### Quantification of HCV RNA by Real-Time Reverse Transcription PCR

HCV RNA in serum and collected livers was extracted using the acid guanidinium-phenol-chloroform method. Quantification of HCV RNA was performed using real-time reverse transcription PCR (RT-PCR) based on TaqMan chemistry, as we described previously (52).

### HCC Analyses

Liver tumors were quantified in fresh livers collected at sacrifice 30 weeks post-HCV infection. Liver tumors were counted, their diameters measured with a ruler and classified as small (0.1 cm diameter, but not exceeding 0.5 cm), medium (0.5–1 cm diameter), or large (>1 cm diameter). Liver tumor burden scores (0–7 scale) were assigned to each liver based on criteria as we described previously (47, 53). The cancerous status of each nodule was verified by qRT-PCR for tumor markers - MCP-1, Timp-1, and glypican-3. The RplP0: Mus musculus ribosomal protein, large, P0 was used as control. The forward primer for MCP-1 was 50-GCATCCACGTGTTGGCTCA-30 and the reverse primer was 50-CTCCAGCCTACTCATTGGGATCA-30. The forward primer for Timp-1 was 50-TGAGCCCTGCTCAGCAAAGA-30 and the reverse primer was 50-GAGGACCTGATCCGTCCACAA-30. The forward primer for glypican-3 was 50-CCAGATCATTG ACAAACTGAAGCA-30 and the reverse primer was 50-CGCAGTCTCCACTTTCAAGTCC-30. The forward primer for 36B4 was 50-TTCCAGGCTTTGGGCATCA-30 and the reverse primer was 50-ATGTTCAGCATGTTCAGCAGTGTG-30. To correct variation in the amount of cDNA available for PCR in the different samples, gene expressions of the target sequence were normalized in relation to the expression of an endogenous control, 36B4 mRNA.

## Results

### HCV infection induces the development of HCC in humanized-liver mice

The main goal of this study is to determine whether the CypI rencofilstat prevents HCV-induced HCC. We first examined whether HCV infection replication from genotypes 1 to 4 (GT1a-GT4a) in humanized mice elicits HCC. As we previously reported (39), HCV replication reaches maximal levels 3 weeks post-infection and remains stable for 30 weeks (Fig 1A). This is true for the 4 genotypes tested. The 4 genotypes induce similar degree of HCC (Fig 1B-D) after 30 weeks.

**Fig 1.**
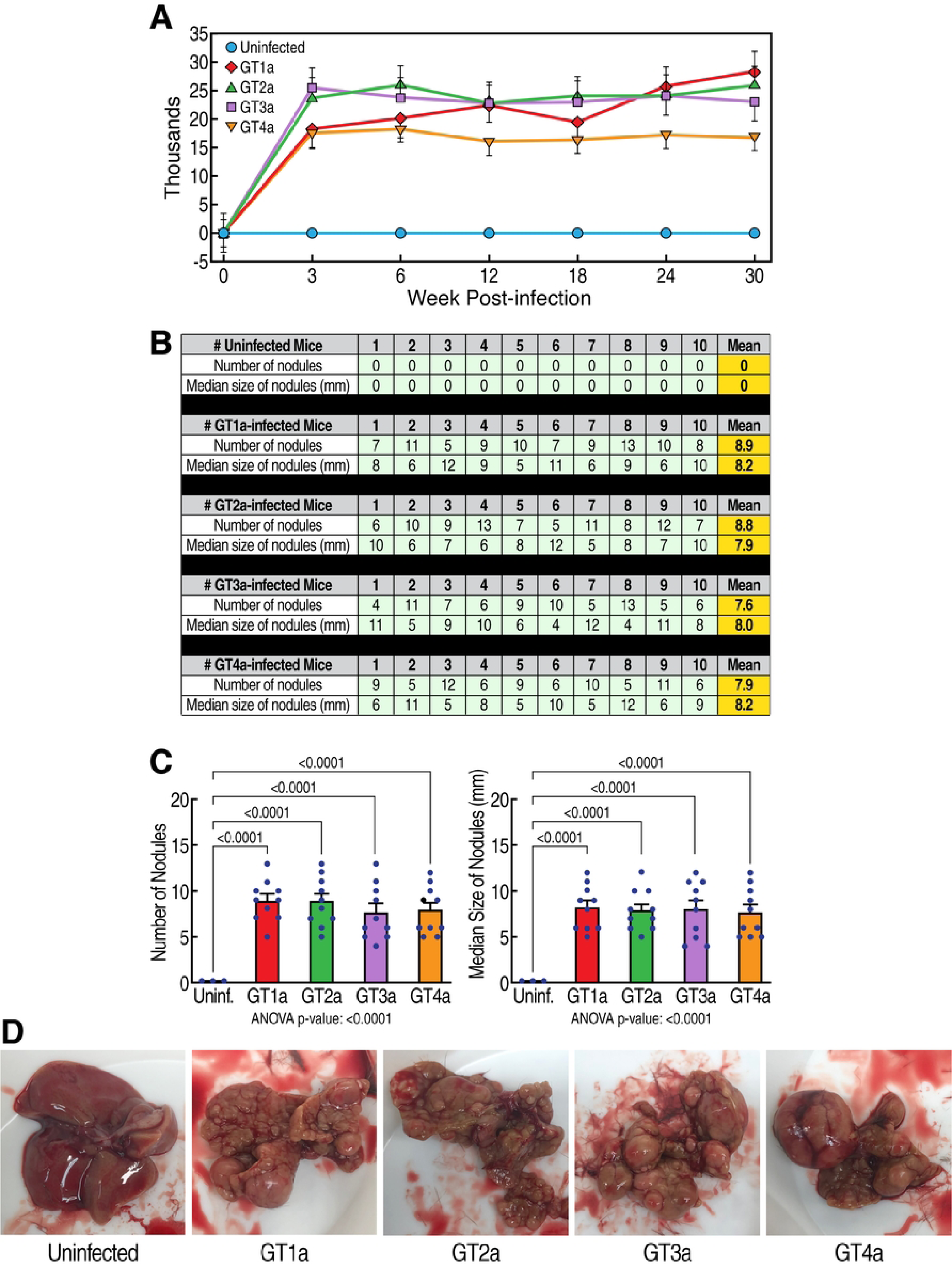
HCV induces HCC. **A.** Humanized MUP-uPA-SCID/Beige mice (n=10) were infected with HCV GT1a, GT2a, GT3a and GT4a and viral infection was monitored by qRT-PCR at the indicated time points. Data are expressed as HCV RNA copies/mL of serum. **B.** Nodules were analyzed in fresh livers collected at sacrifice 30 weeks post-HCV infection. Liver nodules were counted, their diameters measured with a ruler and classified as small (0.1 cm diameter, but not exceeding 0.5 cm), medium (0.5–1 cm diameter), or large (>1 cm diameter). The cancerous status of each nodule was verified by qRT-PCR for tumor markers - MCP-1, Timp-1, and glypican-3 in nodule tissue lysates. **C.** Statistical analyses of B. **D.** Representative liver pictures of uninfected and HCV-infected mice.

### HCV infection induces a progressive HCC development in humanized-liver mice

We next examined the kinetics of the HCV-induced development of HCC. HCV (GT1a)-induced HCC development starts 12 weeks post-infection as demonstrated by the occurence of small cancerous nodules (Fig 2B) and expands over time as demonstrated by higher numbers of small cancerous nodules by 16 weeks (Fig. 2C), and larger cancerous nodules by 24 weeks (Fig. 2D). This demonstrates that HCV infection induces progressive HCC development in humanized MUP-uPA-SCID/Beige mice. These findings are critical because they identified successive stages of HCV-induced HCC progression, and therefore mapped more precisely when rencofilstat treatment could be initiated.

**Fig. 2.**
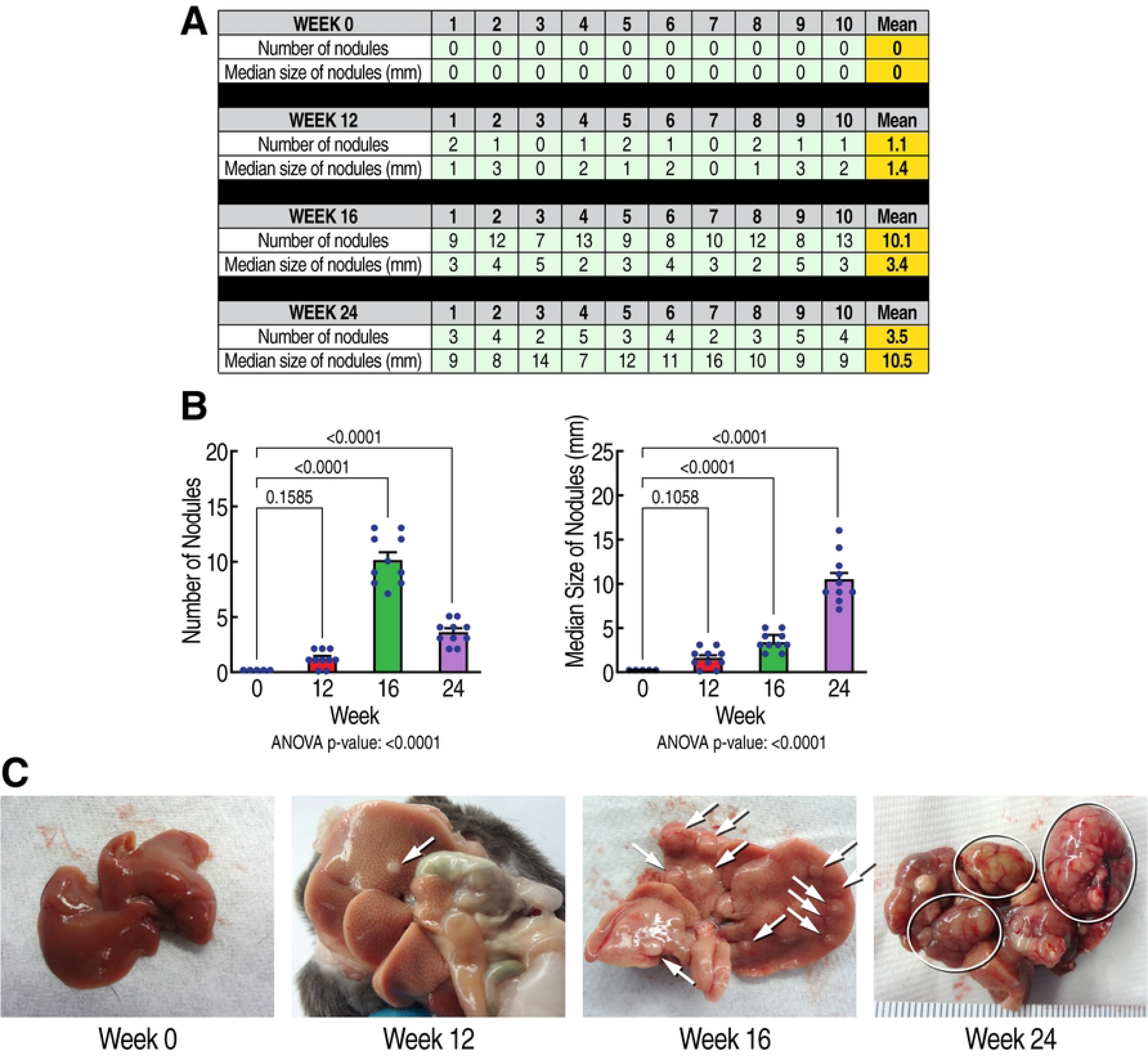
Kinetic of the sequential progression of HCV-induced HCC. **A.** Humanized MUP-uPA-SCID/Beige mice (n=10) were infected with HCV GT1a and liver nodules were analyzed at week 0, 12, 16 and 24 post-HCV infection. Liver nodules were counted, their diameters measured, and their cancerous status was verified by qRT-PCR for tumor markers - MCP-1, Timp-1, and glypican-3. B. Statistical analysis was conducted. **C.** Representative liver pictures of uninfected and HCV GT1a-infected mice.

### Rencofilstat treatment reduces HCV-induced HCC

We next examined the effects of rencofilstat on HCV-induced HCC. GT1a HCV-induced HCC development was induced as described above (Fig. 3, vehicle treatment). Daily rencofilstat treatment initiated at week 0 completely prevented HCC development measured at week 30 (Fig. 3). This was expected since we showed that rencofilstat inhibits HCV infection and therefore eliminated the driver of HCC development (39). In other mice daily rencofilstat administration was initiated when HCC was present in its early, nodular stage (week 12), mid-stage with medium-sized tumors (week 16), or large, late-stage tumors (week 24) as shown in Figure 2. Importantly, daily rencofilstat administration initiated at week 12 not only prevented development of additional nodules but also appeared to regress existing small nodules, since no HCC was observed at week 30 (Fig. 3). More importantly, rencofilstat treatment initiated at week 16, when HCC was well established, significantly decreased tumor sizes at week 30 (Fig. 3). These data indicate that rencofilstat possesses anti-HCC activity, including some degree of regressive activity. This dual beneficial therapeutic effect – anti-HCV and anti-HCC - reveals rencofilstat as a promising anti-HCV-induced HCC therapeutic agent.

**Fig. 3.**
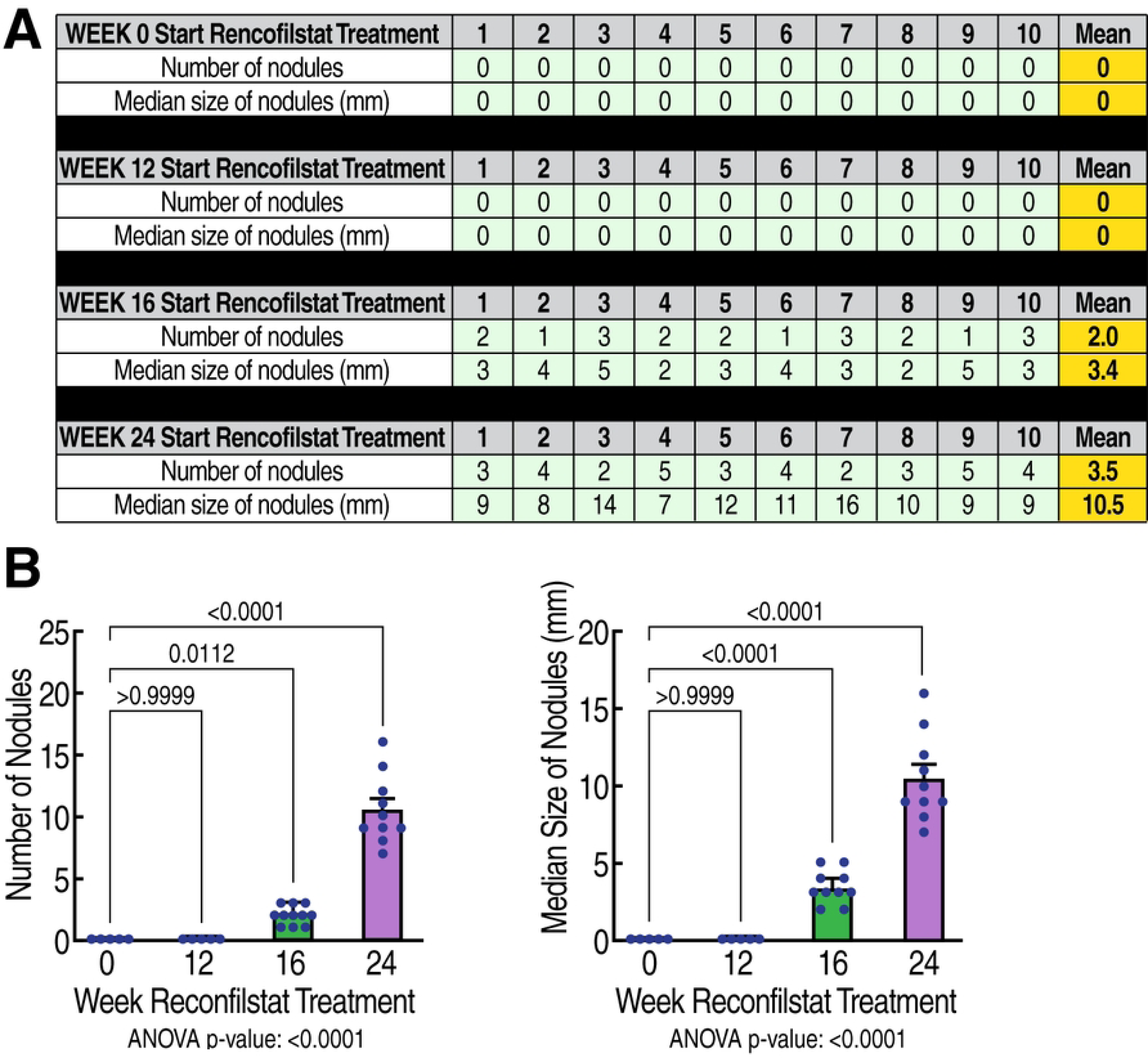
Anti-tumor activity of rencofilstat in HCV-induced HCC. **A.** Humanized MUP-uPA-SCID/Beige mice (n=10) were infected with HCV GT1a and daily treated with rencofilstat starting at week 0, 12, 16 and 24 post-infection. Liver nodules were analyzed at week 24 post-HCV infection. Liver nodules were counted, their diameters were measured and the cancerous status of each nodule was verified by qRT-PCR for tumor markers - MCP-1, Timp-1, and glypican-3. **B.** Statistical analysis was conducted.

### Rencofilstat’s anti-HCC activity is partly independent of its anti-HCV activity

To investigate whether some of rencofilstat’s anti-HCC effects occurs independent of its anti-HCV activity, we compared rencofilstat to other anti-HCV agents in the model. Specifically, we investigated the anti-HCC effect of the NS5Bi sofosbuvir and the NS5Ai inhibitor velpatasvir. As expected, rencofilstat, sofosbuvir and velpatasvir daily treatments starting at 16 weeks post-infection when HCV replication is maximal and when HCV-induced HCC is well established, totally suppressed HCV replication at week 30 (Fig. 4A, last column). Remarkably, rencofilstat, but not sofosbuvir or velpatasvir, also significantly decreased the number of tumor nodules at week 30 by approximately 80% (Fig. 4B). The tumors that remained in the rencofilstat group did, however, grow to the sizes measured in the vehicle, sofosbuvir, and velpatasvir groups. Altogether, the findings demonstrate for the first time that rencofilstat possesses a unique property among anti-HCV agents, that is to exert additional anti-HCC activity independently of its antiviral activity.

**Fig. 4.**
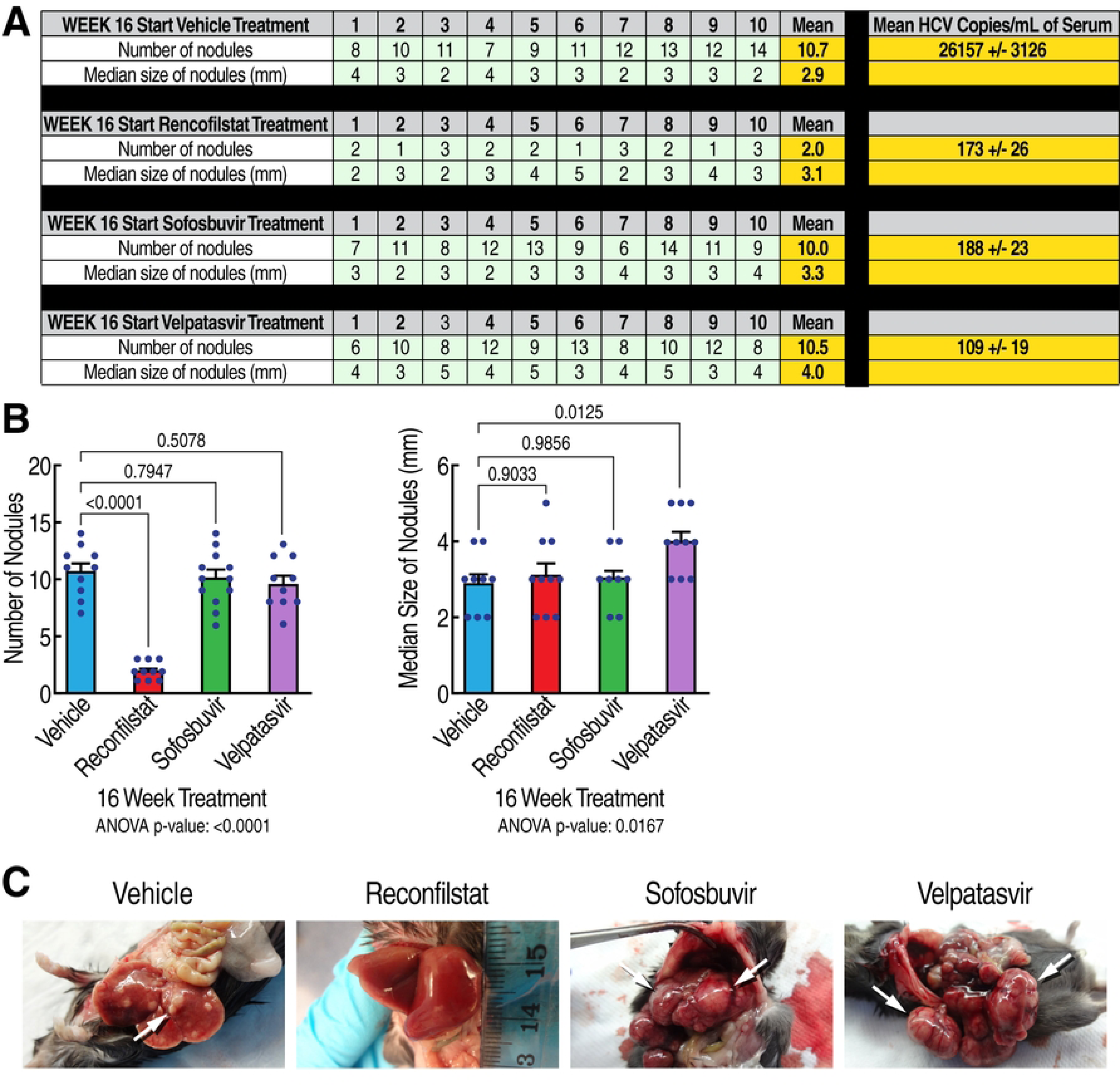
Rencofilstat suppresses HCV-induced HCC mostly independently of its antiviral activity. **A.** Humanized MUP-uPA-SCID/Beige mice (n=10) were infected with HCV GT1a and daily treated with the CypI rencofilstat, the NS5Bi sofosbuvir and the NS5Ai velpatasvir starting at week 16 post-infection. Serum viral loads and liver nodules were analyzed at week 30 post-HCV infection. Liver nodules were counted, their diameters were measured and the cancerous status of each nodule was verified by qRT-PCR for tumor markers - MCP-1, Timp-1, and glypican-3. **B.** Statistical analysis was conducted. **C.** Representative liver pictures of uninfected and HCV GT1a-infected mice.

## Discussion

In the U.S., HCC represents the fastest growing cause of cancer mortality (54–57). HCC accounts for 800,000 deaths per year. Over the past 20 years, the incidence of HCC has more than doubled. Mortality mirrors HCC incidence. An increasing number of young patients have been affected, as the demographic shifts from primarily alcoholic liver disease to those in the 5^th^ to 6^th^ decades of life as the consequences of viral hepatitis acquired earlier in life and in conjunction with high-risk behavior (54–57). In the U.S., the risk factors have historically included alcoholic cirrhosis and viral hepatitis infection. However, the obesity epidemic has resulted in a growing population of patients with nonalcoholic fatty liver disease (NAFLD) and nonalcoholic steatohepatitis (NASH). NAFLD and NASH progress to liver fibrosis, cirrhosis, and HCC. These patients are expected to continuously drive the HCC epidemic worldwide, reflecting the reservoir of the viral hepatitis endemic in the population. Recent studies raised a red flag regarding unexpected higher rates of HCC recurrence following treatment with DAAs for HCV infection (58–71). Studies describing the use of DAAs in HCV patients with HCC are extremely scarce. A recent study assessed the efficacy of DAA regimens in HCV cirrhotic patients who have had HCC compared to those without HCC. Remarkably, 12 weeks post-DAA treatment, patients with liver cancer are 8 times more likely to fail DAA treatment than patients without HCC (72). The authors postulated that HCC serves as a sanctuary for HCV, where viral particles evade DAA therapy (73–74). The increased risk of HCC development following HCV cure is likely the consequence of changes in inflammation (75–77). For example, DAAs rapidly reduce inflammation, but increase serum VEGF levels - a rationale for tumor risk during anti-HCV treatment. Therefore, the identification of drugs, which would interfere with the development of viral hepatitis-induced liver damage, especially HCC, would represent critical tools to enhance the efficacy of DAA treatments. Moreover, the underlying mechanisms of viral reactivation during DAA therapy for HCV are poorly understood. There is thus an urgent need for the identification of new drugs with prolonged effectiveness and with distinct MoA that prevent the development of HCV-induced HCC.

The antiviral MoA of CypI toward HCV is relatively well understood. Specifically, we and others obtained several lines of evidence suggesting that CypI inhibit HCV infection by preventing CypA-HCV NS5A interactions, leading to the prevention of the formation of double membrane vesicles (DMVs), which normally shelter the replication of the viral genome (28). In sharp contrast, the anti-HCC MoA of CypI is poorly understood. In the present study, we showed that rencofilstat inhibited HCV-induced HCC partly and possibly mostly through mechanisms independent of its antiviral activity, suggesting that the CypI rencofilstat possesses anti-cancer properties. Supporting this notion, we reported that rencofilstat inhibits non-viral-induced liver damage including liver fibrosis and HCC as demonstrated by the prevention of cancerous liver nodule formation (14). Several scenarios may explain the beneficial anti-HCC effect of rencofilstat. Since Cyps represent the main targets of CypI, the neutralization of the peptidyl-prolyl isomerase activity of specific members of the Cyp family is likely to be the cause for the anti-HCC activity of rencofilstat. To elucidate which Cyp members participate in HCC development, we created CypA, CypB and CypD knockout (KO) mice, and recently obtained strong evidence that CypD is the main key player in the development of nonviral-induced HCC (manuscript in preparation). Previous work demonstrated that CypD, which resides within mitochondria, may either enhance or decrease tumor growth (78). Since CypD enhances aerobic glycolysis by recruiting hexokinase II to the mitochondrial outer membrane (79–80), it may play a key role in cancer cell metabolism. CypD by inhibiting oxidative stress-induced necrosis may inhibit cell death, and thus enhancing cancer cell survival (81). On the other hand, CypD promotes cancer cell death by enhancing mitochondrial permeability transition pore (mPTP)-mediated apoptosis and necrosis (82–84). These conflicting CypD effects on tumor growth likely result from the broad spectrum of its putative binding ligands that may interfere with mitochondrial permeability. Further studies are required to understand at a molecular and cellular level how CypD regulates HCC development.

## Acknowledgments

We greatly thank Hepion Pharmaceuticals for rencofilstat (CRV431), Dr. D. Geller for fresh human hepatocytes, Dr. A. Kumar for the MUP-uPA-SCID-Beige mice, and Drs. Farci and Lanford for HCV-infected chimpanzee sera. This is publication no. 30246 from the Department of Immunology & Microbiology, The Scripps Research Institute, La Jolla, CA. Research reported in this publication was supported by the National Institute of Allergy and Infectious Diseases of the National Institutes of Health under Award Number R01AI143931. The content is solely the responsibility of the authors and does not necessarily represent the official views of the National Institutes of Health

## Author Contributions

Conceptualization: Philippe A. Gallay.

Formal analysis: Philippe A. Gallay.

Funding acquisition: Philippe A. Gallay.

Investigation: Michael Bobardt, Winston Stauffer, Philippe A. Gallay.

Project administration: Philippe A. Gallay.

Supervision: Philippe A. Gallay.

Validation: Philippe A. Gallay.

Visualization: Philippe A. Gallay.

Writing – original draft: Philippe A. Gallay.

Writing – review & editing: Winston Stauffer, Daren Ure, Robert Foster.

## Notes

### Competing Interest Statement

D. Ure and R. Foster are Hepion Pharmaceuticals employees

